# Ultrasound neuromodulation through nanobubble-actuated sonogenetics

**DOI:** 10.1101/2020.10.21.348870

**Authors:** Xuandi Hou, Zhihai Qiu, Shashwati Kala, Jinghui Guo, Kin Fung Wong, Ting Zhu, Jiejun Zhu, Quanxiang Xian, Minyi Yang, Lei Sun

## Abstract

Ultrasound neuromodulation is a promising new method to manipulate brain activity noninvasively. Here, we detail a neurostimulation scheme using gas-filled nanostructures, gas vesicles (GVs), as actuators for improving the efficacy and precision of ultrasound stimuli. Sonicated primary neurons displayed dose-dependent, repeatable Ca^2+^ responses, closely synced to stimuli, and increased nuclear expression of the activation marker c-Fos only in the presence of GVs but not without. We identified mechanosensitive ion channels as important mediators of this effect, and neurons heterologously expressing the mechanosensitive MscL-G22S channel showed greater activation at lower acoustic pressure. This treatment scheme was also found not to induce significant cytotoxicity, apoptosis or membrane poration in treated cells. Altogether, we demonstrate a simple and effective method to achieve enhanced and more selective ultrasound neurostimulation.

**Graphical abstract:** 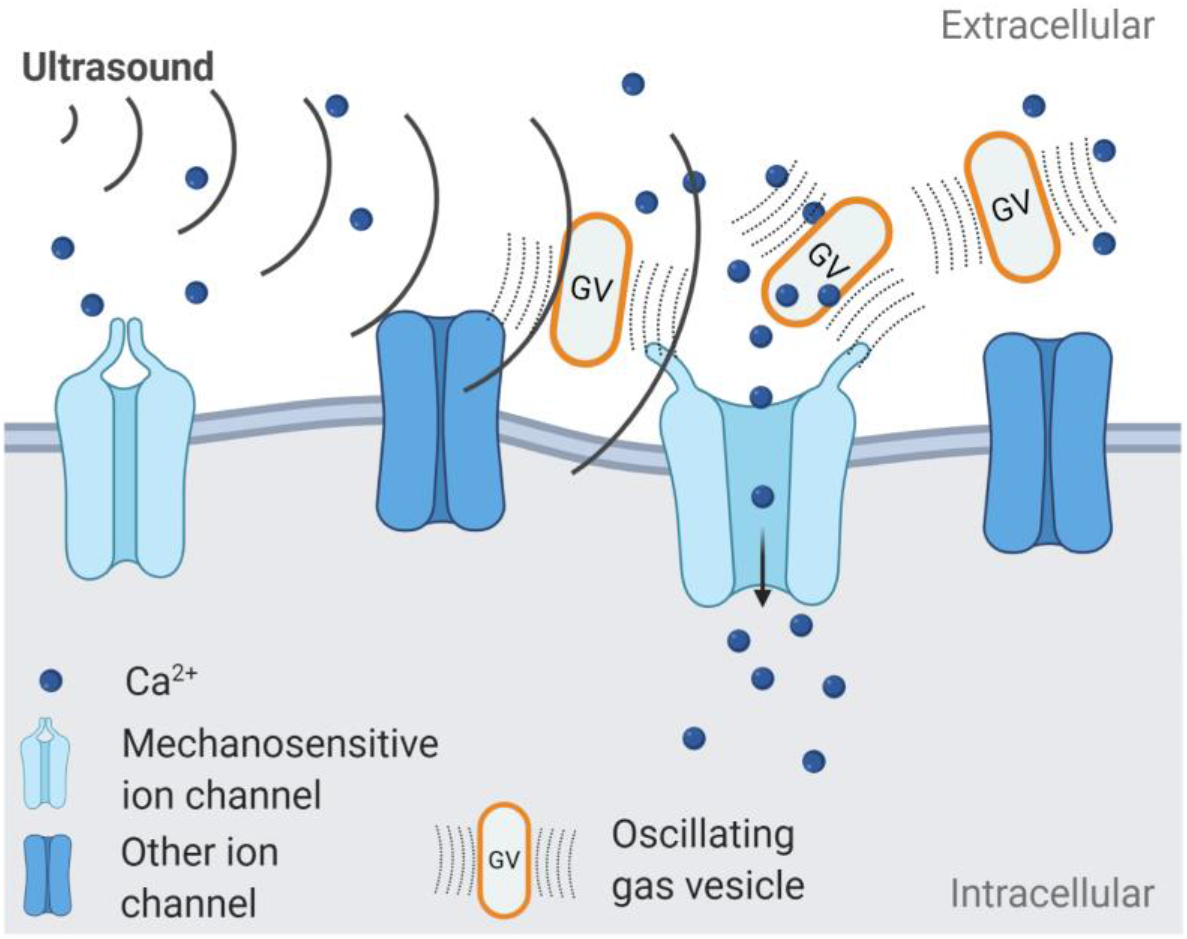

## Introduction

Neuromodulation techniques have expanded greatly over the past decade and have been used to probe neural systems and to treat neurological disorders. Aside from the individual modalities that have been shown to be capable of this, another rapidly-advancing facet of research has been the development of nanoparticles to augment their effects. Such companion nanoparticles have been used as mediators to improve the reach, temporal resolution and targeting of various techniques, or to decrease the invasiveness of the treatment’s scheme. Noteworthy approaches that have been demonstrated include upconversion nanoparticle-mediated (UCNP) near-infrared optogenetics^1^, gold nanoparticle-assisted photothermal stimulation^2^, and magnetic nanoparticle-based magnetothermal/magnetomechanical stimulation^3, 4–5^. Insofar as the goal is to move towards treatments that have high temporal resolution and are as minimally-invasive as achievable, nanoparticle mediators have helped in this regard.

Ultrasound is mechanical energy, able to non-invasively target deep-seated regions in the brain and be focused to spots a few millimeters size. It is also an emerging neuromodulation technique which can be used to modulate brain activity safely^6, 7^ Experiments in many different animal species showed successful stimulation of various brain regions, such as rodents^8^, rabbits^9^, pigs^10^, sheep^11^, non-human primates^12^. Low-intensity ultrasound has been used to stimulate various brain regions in the human, including the thalamus^13^, the prefrontal, visual^14^, motor^15^ and somatosensory^16, 17, 18, 19^ cortices. It is also under study as a possible treatment for a range of neurological disorders, such as Alzheimer’s disease^16, 17, 18, 19^, Parkinson’s disease^20, 21^, epilepsy^22^, depression^23^ and amyotrophic lateral sclerosis^24^. Ultrasound has thus shown the ability to affect the functioning of the central nervous system without significant accompanying thermal damage. One mechanism through which ultrasound is understood to exert such effects is by activating mechanosensitive ion channels present in the cell membranes, whether endogenous or externally-introduced^25, 26, 25, 26^. There is also understood to be some endogenous level of mechanosensitivity in most cells, including those of the brain^27^. In light of this, it would also be helpful to be able to limit the effects of ultrasound to a desired area and cell-type which could help minimize side-effects and increase treatment efficacy. It would, hence, be useful to be able to increase the targetability of ultrasound stimulation.

A candidate for such a tool is nano-sized bubbles extracted from cyanobacteria, called gas vesicles (GVs). GVs produce robust ultrasound contrast signals and have been applied to serve as reporters for specific genes, molecules and cellular activities^28^. The non-linear signals thus generated come from ultrasound driven buckling effects which are frequency-independent. The oscillations can also generate mechanical perturbations to the surrounding environment^29^. Given these unique acoustic properties, we hypothesized that GVs could oscillate in a low-frequency ultrasound field and serve as actuators to activate mechanosensitive ion channels to induce neurostimulation effects.

In the present study we demonstrate a GV-actuated strategy to achieve controllable ultrasound neuronal stimulation. We were able to stimulate primary neurons with low intensity ultrasound in the presence of GVs to induce Ca^2+^ influx, but not otherwise. The neuronal responses were dose-dependent and reversible, as well as closely temporally-tied to the ultrasound stimuli. The combination of GVs and ultrasound (referred to as ‘GVs+US’) also significantly increased the expression of c-Fos in the nuclei of neurons, a further indication of activation. We also show that the stimulation effect was mediated in large part by the activation of mechanosensitive ion channels. Building on this finding, we induced heterologous expression of a mechanosensitive ion channel in neurons, and we were able to reduce the acoustic intensity level required to induce calcium response and neuronal c-Fos expression. The stimulatory effects were also found to be mostly limited to the neurons expressing the channels. The combination of US+GVs was safe and non-harmful to the treated cells. Thus, we provide evidence for an enhanced ultrasound *in vitro* neuromodulation method capable of increasing response to ultrasound and detail how the treatment may be made more targeted through the mechanism of mechanosensitive ion channels.

## Results

### Characterization of GVs’ properties

GVs were prepared from *Anabaena flos-aquae* through tonic cell lysis and centrifugally-assisted flotation30. They were found to typically be 50 - 100 nm in width and 100 - 500 nm long (Fig. 1A-B). The zeta potential of GVs was −40 ± 5 mV, indicating a suitable surface charge for colloidal stability (Fig. 1C)^31^. We also found that our prepared GVs were not cytotoxic on their own to primary neurons in culture (Fig. 1D). In all, our prepared GVs were found to be nano-sized, stable and non-cytotoxic.

**Fig. 1.**
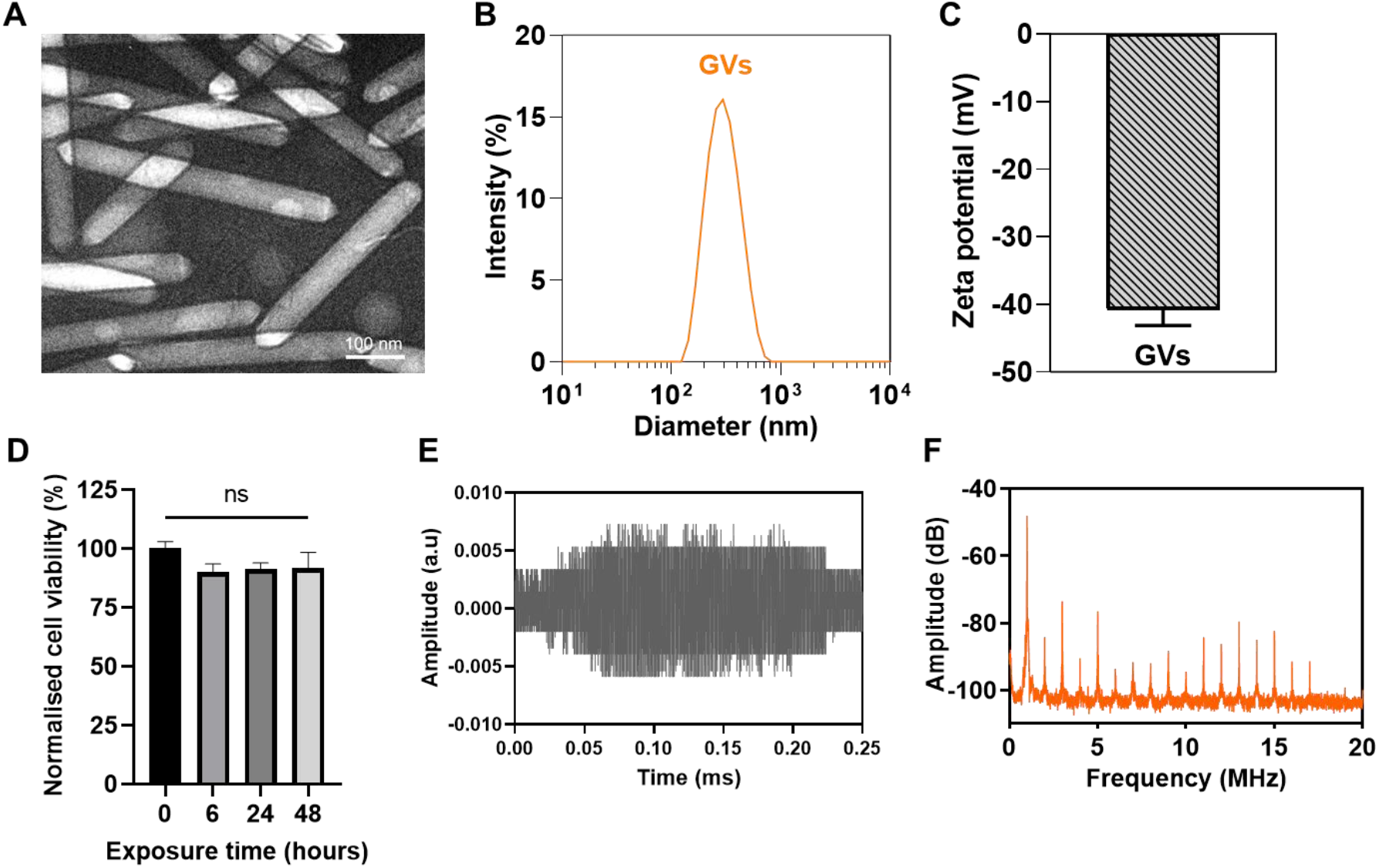
Basic characterization of the prepared GVs. **a,** Transmission electron microscopy (TEM) image of the prepared GVs. Scale bar represents 100 nm. **b,** Number-averaged diameter of GVs in deionized (DI) H_2_O as measured by Dynamic Light Scattering (DLS). Data represent the mean of 3 independent experiments. **c,** Zeta potential of GVs in DI H_2_O. Bar represents mean ± SD of 3 independent experiments. **d,** Cytotoxicity of GVs (0.8 nM), as measured by an MTT test. Primary neurons were exposed to GVs in medium for the stated amounts of time. Bars represent the mean ± SEM of 3 independent experiments. No significant differences were found by one-way ANOVA. **e,** Representative time-domain waveform of backscattered signals from a purified GV suspension (0.8 nM) sonicated by a 1.0 MHz tone burst sinusoidal wave at 0.28 MPa PNP, after one burst interval (300 cycles). **f,** Averaged frequency spectrum of backscattered signals from purified GVs suspension under the same sonicating conditions as in **(e)**.

Ultrasound is known to induce both stable and inertial cavitation^32^. Generally, stable cavitation occurs at relatively low ultrasound intensities, caused by size changes of gas-filled bubbles in a sustained, periodic manner. Inertial cavitation usually occurs at high ultrasound intensities, when gas bubbles collapse, generating a shock wave that could cause significant cell damage. We wanted to control ultrasound intensity such that it would enable the GVs to generate robust stable cavitation but not inertial cavitation, which required characterizing the GVs’ responses in an ultrasound field. Hence, we performed passive cavitation detection using a setup in a tank of degassed water, where GVs suspensions were exposed to pulsed ultrasound at 0.28 MPa peak negative pressure (PNP) (schematically illustrated in Supplementary Fig. 1A). We observed the backscattered signals in the time-and frequency-domains to monitor the patterns of cavitation produced. We found no broadband signal and only the appearance of 1^st^ - 17^th^ harmonic signals (Fig. 1D-E), indicating that no inertial cavitation occurred when the GVs were sonicated in our setup. Crucially, 0.28 MPa was the highest acoustic pressure used in the entire study, making inertial cavitation unlikely at the range of intensities used in the various following experiments.

### Customized *in vitro* ultrasound stimulation setup

For the present study, we used a customized system which facilitated ultrasound stimulation and calcium imaging simultaneously (Fig. 2A). Briefly, the ultrasound stimulation system was aligned with a calcium imaging system and the calcium responses of the stimulated neurons were monitored. Ultrasound was delivered through a waveguide filled with degassed water that was attached to the ultrasound transducer assembly. Cells were cultured on glass coverslips placed inside a culture dish, and GVs were added to the medium and gently mixed just before stimulation. Prior to cellular stimulation, we tested the acoustic pressure and field produced by this setup using a hydrophone, and found that it provided a relative homogeneous ultrasound field in the central region (Supplementary Fig. 1B). Each stimulus was composed of 300 tone burst pulses at a center frequency of 1.0 MHz, 10% duty cycle, pulse repetition frequency (PRF) of 1 kHz, at low acoustic intensities (0.03 - 0.20 MPa). These parameters amounted to ultrasound being delivered in very short bursts, minimize thermal effects. For experiments not involving real-time imaging, cells were treated inside a standard cell culture incubator, as described in our previous study^33^. Parameters for stimulation in these experiments was the same as mentioned above, with a slightly different range of acoustic pressures (0.14 - 0.28 MPa) and a treatment time of 15 minutes.

**Fig. 2.**
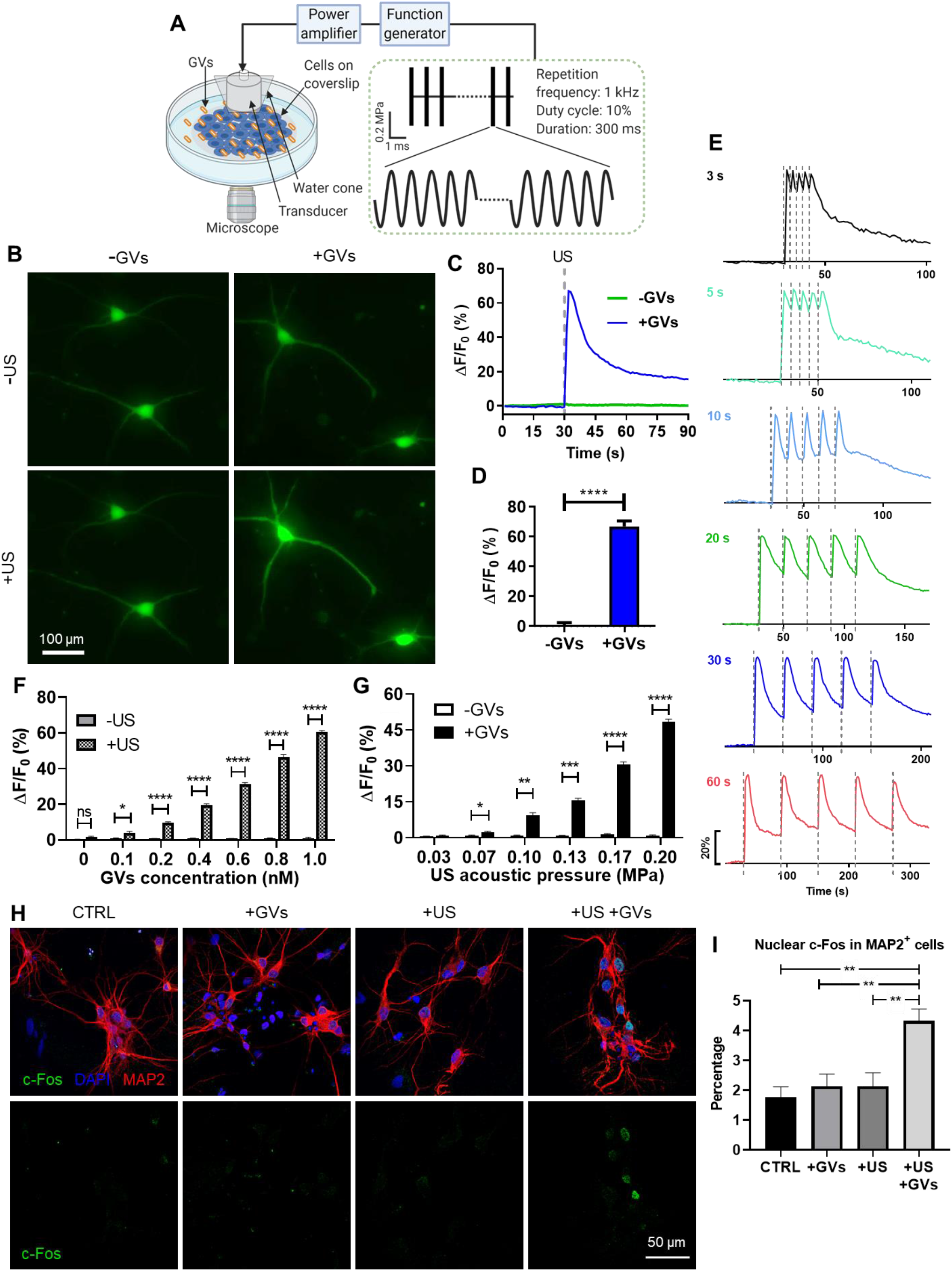
GVs enable low-intensity ultrasound to stimulate activity in primary neurons. **a,** Schematic illustration of the GV-mediated ultrasound setup for recording cells. GVs were mixed into cell culture medium. Cellular response upon US+GVs stimulation was observed in real time. **b,** Representative images of GCaMP6s fluorescence in primary neurons with or without GVs, before and after 0.20 MPa ultrasound. **c,** Ca^2+^ imaging time course of neurons in (**b**). **d,** Ca^2+^ response of neurons to stimulation by 0.20 MPa ultrasound. Bars represent mean ± SD from 3 independent experiments. ****, p < 0.0001, two-tailed unpaired *t*-test. **e,** Time-resolved Ca^2+^ responses of neurons stimulated by 5 ultrasound pulses at varying intervals. **f,** Ca^2+^ response of cells to varying ultrasound intensities, 0,8 nM GVs. Bars represent mean ± SEM of 3 independent experiments. *, p < 0.05, **; p < 0.01; ***, p < 0.001; ****, p < 0.0001, two-tailed unpaired *t*-test with Holm-Sidak correction. **g,** Ca^2+^ response of cells to varying GV concentrations, 0.20 MPa ultrasound. Bars represent mean ± SEM of 3 independent experiments. *, p < 0.05; p < 0.0001, two-tailed unpaired *t*-test with Holm-Sidak correction. **h,** Representative IF images of c-Fos and MAP2 staining in untreated cells, and cells treated with the indicated combinations of ultrasound (0.28 MPa) or GVs (0.8 nM). **i,** Quantified results of nuclear c-Fos staining in MAP2^+^ cells after various treatments, as in **(h)**. Bars represent mean ± SEM from 4 independent experiments. **, p < 0.01, one-way ANOVA with post-hoc Tukey test.

### GVs enable efficient neuromodulation by low-intensity ultrasound

We first observed the Ca^2+^ response in rat primary cortical neurons when stimulated with ultrasound and GVs (1.0 MHz center frequency, 0.20 MPa ultrasound and 0.8 nM GVs unless otherwise indicated). Neurons were made to express the genetically-encoded calcium sensor GCaMP6s under the human synapsin promoter using AAVs, and we monitored its fluorescence upon stimulation. We found that GCaMP6s fluorescence increased quickly and dramatically when stimulated with an ultrasound pulse in the presence of GVs, but not without them, and the fluorescence gradually returned to the baseline without further stimulation (Fig. 2B-D). Next we tested whether neuronal activation could be induced repeatedly. Five 300 ms pulses were delivered to cells at intervals of 3, 5, 10, 20, 30 or 60 seconds in the presence of GVs and the temporal profiles of the cells’ Ca^2+^ response was charted. Stable and reversible calcium transients were seen to quickly follow each pulse, and the neurons were able to recover after each pulse when given enough time (max ΔF/F = 46 ± 1.8%, five pulses) (Fig. 2E & Supplementary Video 1). Aside from primary neurons, we also observed the same pattern of responses, albeit at lower amplitudes, in the mouse hippocampal cell line mHippoE-18 (referred to as ‘CLU199’ in this manuscript). In the presence of GVs, ultrasound triggered robust, repeatable, and rapid calcium responses from cells after which calcium levels would gradually recover, while no response was seen without GVs (Supplementary Fig. 2A-D). Furthermore, the cellular response was found to vary with, both, the concentration of GVs and the acoustic pressure applied, both in primary neurons (Fig. 2F-G) and in CLU199 cells (Supplementary Fig. 2E-F). These data help to establish that the responses observed were indeed caused by the US+GVs treatment, and also reveal how such a combination treatment can easily be tweaked to suit the degree of response desired. A GVs concentration as low as 0.1 nM was sufficient to induce a significantly higher response than without GVs, showing that GVs could indeed lower the acoustic pressure threshold needed for Ca^2+^ response, which would otherwise require increased ultrasound intensity^7^. Finally, when treated with US+GVs primary neurons showed approximately double the nuclear c-Fos expression, a marker of neuronal activity downstream of Ca^2+^ influx^34^, than when they were untreated or exposed to only GVs or ultrasound (Fig. 2H-I). Thus we were able to use GVs to efficiently activate primary neurons with short bursts of low-intensity ultrasound.

### GV-mediated ultrasound stimulates cells by activating mechanosensitive ion channels in the cell membrane

A possible confounding factor in our experiments was that sonoporation, of the kind that is typically induced by ultrasound in the presence of microbubbles, is known to play a role in initiating Ca^2+^ response^37^, 38. Although only stable cavitation was detected in our system, that evidence could not address whether our treatment was causing pore formation in the membrane. Thus, we performed a membrane integrity assay to see whether sonoporation was involved in the Ca^2+^ responses to the GV-mediated ultrasound treatment. We used the membrane impermeable dye propidium iodide (PI) and observed whether it could penetrate the cell membrane during the stimulation. Insonated primary neurons in the presence of GVs evoked Ca^2+^ influx but no PI could be detected inside the cells; brightfield imaging also showed that the cells maintained their morphology following the treatment (Fig. 3A, Supplementary Fig. 3A & Supplementary Video 3). To contrast, Triton X-100 was used as a positive control for membrane permeation^35^, and PI influx was seen within 30 seconds of its addition and continued to increase for the remainder of the assay, while the intracellular calcium signal decreased and the cell was visibly damaged (Fig. 3A & Supplementary Fig. 3A). In general, neither ultrasound alone nor the +GVs condition used in our experiments were seen to trigger PI influx in primary neurons or CLU199 cells (Supplementary Fig. 3B-C). Further, we did not observe obvious cytotoxicity or apoptosis in primary neurons following the treatments (Supplementary Fig. 3 D-E). Thus, we concluded that our US+GVs treatment could trigger calcium responses in cells with negligible loss of membrane integrity, which is consistent with stable cavitation hypothesis.

**Fig. 3.**
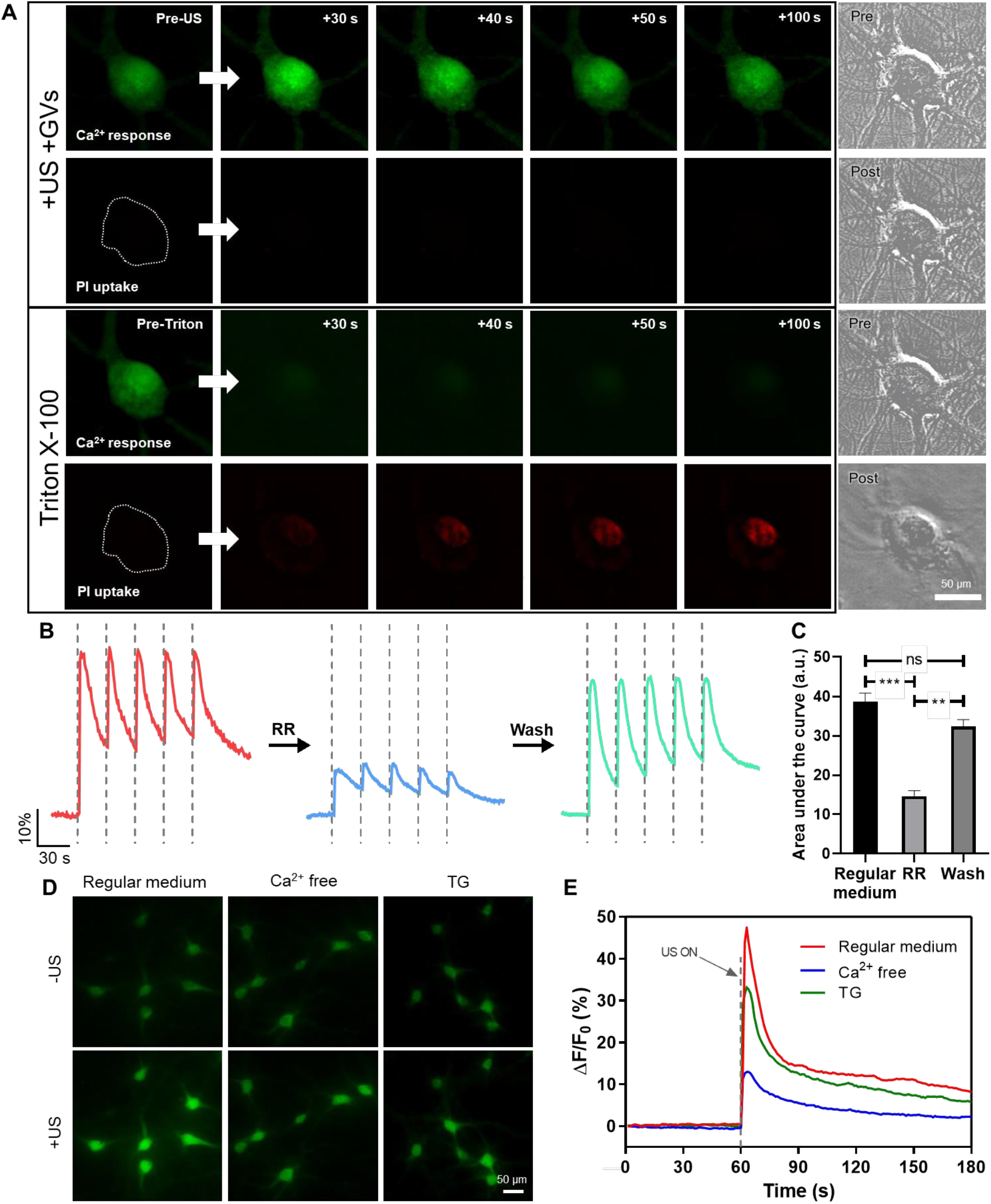
Mechanosensitive ion channels are an important mechanism for GV-mediated ultrasound stimulation. **a,** Upper: Calcium response and PI uptake (indicating membrane integrity) during US+GVs stimulation. Lower: Calcium response and PI uptake (indicating membrane integrity) following addition of 0.2 mM Triton X-100 as a positive control for loss of membrane integrity. Right: Brightfield images of the images cells before and after ultrasound + GVs or Triton X-100. All images shown in this panel are representative. **b,** Time-resolved calcium responses of neurons during US+GVs stimulation, first as normal, then in the presence of mechanosensitive ion channel blocker ruthenium red (RR), and then after RR was washed away. **c,** Quantification of area under the curve before, during and after RR treatment as shown in **(b)**. Bars represents the mean ± SEM of 3 independent experiments. **p < 0.01, ***p < 0.001, one-way ANOVA with post-hoc Tukey test. **d,** Representative images of calcium responses of neurons in regular medium, Ca^2+^-free solution and Thapsigargin (TG). **e,** Time-course calcium imaging of cells before and after ultrasound + GVs stimulation in the three solutions indicated in **(d)**.

We next tried to address whether calcium influx by mechanosensitive ion channels was in fact responsible for the observed effects of the treatment. We used Ruthenium Red (RR, 20 μM), a blocker for a wide range of mechanosensitive ion channels^36^, to see if the response to US+GVs would be altered. Calcium responses to ultrasound pulses were found to be significantly suppressed in the presence of RR, and the responses recovered when it was washed away (Fig. 3C-D). We observed a similar suppression of Ca^2+^ influx in the presence of RR in CLU199 cells (Supplementary Fig. 2A-C). We then tried to identify the main source of Ca^2+^ responses by treating cells with US+GVs in EGTA-chelated medium, or pre-treating cells with Thapsigargin (TG, 3 μM) to deplete intracellular calcium stores37. Compared to normal conditions, cells in Ca^2+^-free medium showed much reduced calcium influx (~75% reduction), but the reduction in TG-treated cells was much lesser (~ 25-30% reduction) (Fig. 3D-E & Supplementary Fig. 2A-C). We thus found that while intracellular Ca^2+^ release played some role in the US+GVs response, calcium influx from the external medium had a much larger contribution to the observed outcomes. Putting our evidence together, with no significant sonoporation being observed, cellular response being significantly depressed when treated with RR or in Ca^2+^-free medium but not in TG-treated cells, we inferred that activation of mechanosensitive ion channels was an important mechanism of GV-mediated ultrasound stimulation.

### Increased levels of mechanosensitive ion channels improve selectivity of GV-mediated ultrasound stimulation

Having seen the important role mechanosensitive ion channels play in GV-mediated ultrasound, we surmised that one way of increasing the sensitivity and efficiency of our treatment would be to increase expression of mechanosensitive ion channels in desired cells. Furthermore, since methods of inducing expression of a desired protein (such as AAVs) offer the option of cell-type selectivity, we hypothesized that we could also improve the targeting of the stimulation to neurons. Our general idea was to induce the expression of a mechanosensitive ion channel in primary neurons, which would sensitize them to GV-mediated ultrasound stimulation at lowered intensities (illustrated schematically in Fig. 4A).

We chose a mutant version of the well-studied bacterial channel large conductance mechanosensitive ion channel (MscL-G22S)^42^. In our previous work, we induced neurons in certain mouse brain region to express this channel and found that we were able to both sensitize neurons to ultrasound alone, and target brain regions using this method^38^. We used human synapsin-promoted AAVs encoding for MscL-G22S-EYFP (called ‘MscL-EYFP’), or EYFP alone as a control, to transduce primary neurons. EYFP fluorescence was used to identify cells that were transduced successfully, as described in our previous study^38^. To evaluate the ability of MscL to sensitize neurons to acoustic radiation force, we treated them with lowered acoustic intensities (0.07 to 0.17 MPa). For calcium imaging experiments, the GV concentration was also halved to 0.4 nM to more clearly observe the effects of greater neuronal mechanosensitivity without saturating our imaging system. We found that the MscL+US+GVs condition showed a rapid and strong response to one pulse of 0.13 MPa ultrasound, with all other conditions showing much lower or no response (Fig. 4B & Supplementary Video 3). Crucially, we found that cells expressing MscL-EYFP showed a significantly higher response to ultrasound stimulation than did cells without EYFP in the same dish (Supplementary Fig. 4A). MscL+US+GVs also showed the strongest response to 3 out of 4 ultrasound intensities tested, and significantly higher response than EYFP+US+GVs at the 2 intermediate intensities (Fig. 4C). The lowest ultrasound intensity at which a differential response between the +GVs and-GVs groups could be observed was 0.10 MPa. The EYFP group did not show obvious responses at any of the tested intensities, while the MscL and EYFP+GVs groups showed responses with increasing ultrasound intensity, indicating the sensitizing role of both MscL expression and GVs. Interestingly, as in our previous study, we found again that as ultrasound intensity is increased, the strength of the EYFP and MscL-EYFP conditions’ responses converge until they do not differ significantly^38^. We also found that when stimulated by 0.14 MPa ultrasound and 0.8 nM GVs, MscL+US+GVs neurons showed significantly higher nuclear c-Fos expression than all other groups tested, including EYFP+GVs (Fig. 4D-E & Supplementary Fig. 4B). We also compared the cells expressing MscL-EYFP vs those that did not in the same dish, and found that the EYFP^+^ cells showed significantly higher nuclear c-Fos levels (Fig. 4F). Thus, we demonstrate that increasing the expression of mechanosensitive ion channels in desired cells can increase their sensitivity to ultrasound, and that their preferential expression in specific cell-types is an effective way to target the effects of GV-mediated ultrasound stimulation.

**Fig. 4.**
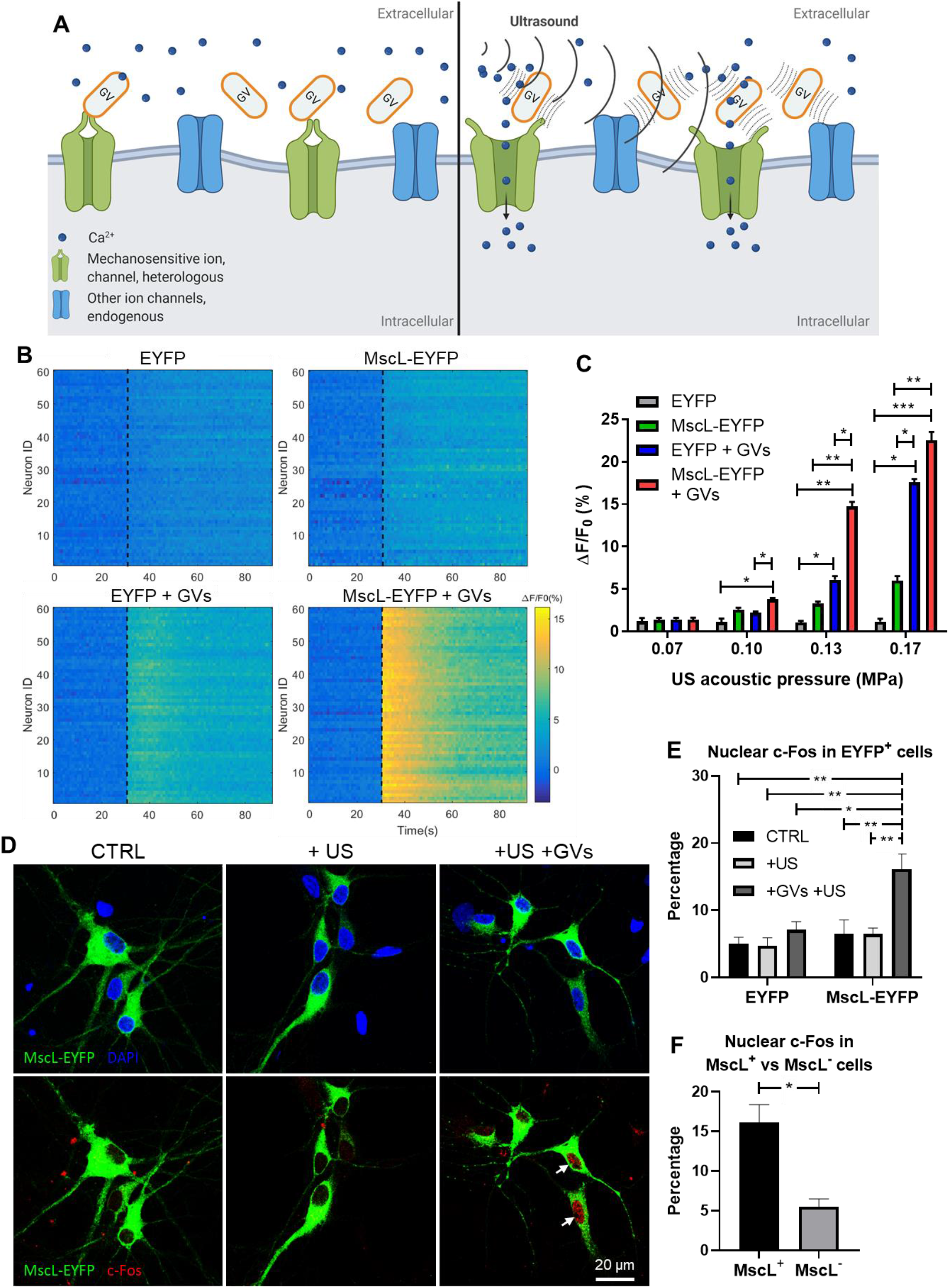
Increased expression of a mechanosensitive ion channel increases neurons’ sensitivity to GV- mediated ultrasound stimulation. **a,** Schematic illustration of the sonogenetic experimental scheme. Briefly, a heterologous mechanosensitive channel, MscL-G22S is expressed preferentially in primary neurons, which increases their sensitivity to ultrasound + GVs treatment, enabling cell-type specific activation. **b,** Temporal raster plots of fluorescence changes in neurons, transducted with EYFP or MscL-G22S-EYFP AAVs, stimulated by a single 300 ms ultrasound pulse (0.13 MPa, black dashed line) with or without GVs. **c,** Calcium responses in AAV-transducted neurons after ultrasound stimulation with varying acoustic pressures, with or without GVs. Bar chart represents means ± SEM of 3 independent experiments. *p < 0.05; **p < 0.01; ***p < 0.001; ****p < 0.0001, two-way ANOVA with post-hoc Tukey test. **d,** Representative images of neuronal c-Fos expression in cells transducted with MscL-EYFP in untreated cells, or cells treated with ultrasound alone or ultrasound + GVs. **e,** Quantified results of nuclear c-Fos staining in MscL-EYFP^+^ cells after treatments, as in **(d)**. Bars represent mean ± SEM from 3 independent experiments. **, p < 0.01; one-way ANOVA with post-hoc Tukey test. **f,** Quantified results of nuclear c-Fos staining in cells with and without MscL-EYFP in the same dish, following ultrasound + GVs stimulation. Bars represent means ± SEM of 3 independent experiments. *, p < 0.05, two-tailed unpaired *t*-test.

## Discussion

Non-invasive neuromodulation technologies with cell-type selectivity hold great potential for studying neural circuits and treating neurological conditions. Here we present a GV-mediated sonogenetic toolkit for selective neuronal activation, comparable to previously-demonstrated magnetothermal and magnetomechanical approaches. We employed a 1.0 MHz transducer and demonstrated that GVs can serve as localized acoustic amplifiers to decrease the threshold of ultrasound intensity for neurostimulation. We achieved controllable calcium signaling in both primary neurons and a neuronal cell line, and increased c-Fos expression in primary neurons with low-intensity ultrasound stimulation in the presence of GVs through activation of endogenously-expressed mechanosensitive ion channels. We also showed that combining increased mechanosensitivity (through expression of MscL in neurons) with GVs can significantly increase cells’ ability to respond to lowered acoustic pressures, and that the stimulation can potentially be limited to the targeted cells. These data suggest that the GV- mediated ultrasound reliably and reversibly activates cells *in vitro* by opening mechanosensitive ion channels, and that the treatment is generally not harmful to cells.

A limitation of this study is that it lacks information about ion-channel dynamics, which could be obtained by patch clamping during ultrasound stimulation. We found this to be unachievable during the present study as we could not eliminate vibration of the patch pipettes when ultrasound was turned on, which has been reported in other studies as well^39, 40^. Indeed, it would be useful to know the specific parameters required to trigger channel opening in GV-mediated ultrasound stimulation. However, we were able to collect evidence of the role of mechanosensitive ion channels through calcium imaging by using various blockers and calcium-free imaging. We also showed that the cells’ calcium influx through ion channels response was independent of, and indeed greater than, intracellular calcium release or calcium influx through sonoporation. The expression of MscL in neurons was seen to be an effective method of increasing cells’ sensitivity to ultrasound, even when the concentration of GVs was lowered.

We also showed that cells expressing MscL showed significantly higher calcium response and nuclear c-Fos than other cells in the same dish that did not, which indicates that US+GVs stimulation does act on mechanosensitive ion channels. Thus, we provide alternative data for the activation of mechanosensitive ion channels by tracking effects downstream of channel opening.

Consistent with our previous study, we found the expression of the MscL channel in neurons to be effective in increasing cellular sensitivity to ultrasound^38^. We detected low Ca^2+^ response from cells without MscL, but obvious and significantly greater responses from MscL-expressing cells when stimulated with low-intensity US+GVs. The observed MscL-dependence of these effects in our experiments pinpoints the channel as the key source of improved sensitivity to stimulation. The performance of this approach may be further improved by using other ultrasound-sensitive ion channels or using different or novel mutants of the MscL channel. By the same logic, our scheme may be applied to identify natural problematic situations in which cells show increased expression of mechanosensitive ion channels, such as in aging brains^41^ or in the progression of some cancers or infectious diseases^42, 43^. Thus, the ability of GV-mediated ultrasound stimulation to act upon mechanosensitive ion channels could be applied in fields beyond neurostimulation.

A recent study from our group demonstrated that surface-modified GVs can escape clearance from the reticuloendothelial system and penetrate tumor vasculature through enhanced permeability and retention (EPR) effects^44^. In addition, expressing GVs as acoustic reporter genes (ARGs) in mammalian cells to enable ultrasound imaging of mammalian gene expression have been reported^45^. Such application of ARGs could be a milestone development for ultrasound imaging, almost analogous to the role of green fluorescent protein (GFP) in optical imaging. Alternatively, GVs can also be delivered to targeted regions, since it is nano-sized. Microbubble-mediated ultrasonic bio-effects have been widely explored and utilized to open cell membranes and the blood-brain barrier^46, 47, 48, 49, 50^. However, the micrometer size of microbubbles limits spatial resolution and they are restricted to use in blood vessels due to their size and have a short half-life in vivo (<5 min in the blood)^51, 52^. In contrast, GVs are gas-filled protein-shelled nanostructures and these protein shells exclude water but permit gas to freely diffuse in and out from the surrounding media, making them physically stable despite their nanometer size. Therefore, US+GVs could even be developed to have a more theranostic role in the brain.

## Acknowledgements

This work was supported by the Hong Kong Research Grants Council General Research Fund (15102417 and 15326416), Hong Kong Innovation Technology Fund Mid-stream Research Program (MRP/018/18X), Key-Area Research and Development Program of Guangdong Province (2018B030331001), and internal funding from the Hong Kong Polytechnic University (1-ZE1K and 1- BBAU). The authors would like to thank the facility and technical support from University Research Facility in Life Sciences (ULS) of The Hong Kong Polytechnic University. Schematic illustrations seen in the graphical abstract, Fig. 2A and Fig. 4A were created with BioRender.com

## Author contributions

Conceptualization, X.H., Z.Q. and L.S.; Methodology, X.H., Z.Q., S.K., J.G. and L.S.; Investigation, X.H., S.K., K.F.W., T.Z. and M.Y.; Data Analysis, X.H., Z.Q., S.K., J.Z. and Q.X..; Manuscript Preparation, H.X, Q.Z., S.K. and L.S.; Supervision, J.G. and L.S.; Funding Acquisition, L.S.

## Declaration of interests

The authors have submitted a patent application titled “A non-invasive method for selective neural stimulation by ultrasound” with the U.S. Patent and Trade Office, dated April 10, 2018, assigned application number 15/949,991. The authors declare no further financial interests.

## Materials and methods

### Gas vesicle preparation

*Anabaena flos-aquae* was cultured in sterile BG-11 medium at 25 °C under fluorescent lighting with 14 hours/10 hours light/dark cycle. GVs were isolated by hypertonic lysis to release GVs by quickly adding sucrose solution to a final concentration of 25%. GVs were isolated by centrifugation at 400 x *g* for 3 hours after lysis. To purify GVs, the solution was washed by the same centrifugation process 3 times and stored in PBS at 4 °C. The GVs’ concentration was measured by optical density at 500 nm (OD500) by UV-Visible spectrophotometer^32^.

### Passive cavitation detection

Acoustic spectroscopy on GV suspensions were performed in a custom-built chamber, the 1 MHz flat transducer and hydrophone (HGL-0200, Onda) were perpendicularly aligned and immersed in a tank of deionized, degassed water (Fig S1A). A rectangular agarose (3%) chamber of wall thickness 5 mm and cavity 15×15 mm was placed in the middle, with the center point 17.5 mm away from both the transducer and the hydrophone. 1 MHz sinusoidal trains of burst width 200 μs and burst interval 2 ms were generated by a function generator (AFG251, Tektronix), amplified by a radio frequency (RF) amplifier (A075, Electronics & Innovation Ltd.), to drive the emitting transducer, producing acoustic output with 0.28 MPa peak negative pressure. Signals received by the hydrophone were amplified (AH- 2010, Onda) and digitized (CSE1222, GaGe) before analysis. 20 sections of 200 μs digitized signal in 20 separate bursts were processed with fast Fourier transform (FFT) using MATLAB and the resulting frequency spectra were averaged.

### Cell culture

All cells were grown inside a standard humidified cell culture incubator at 37 °C with 5% CO_2_. CLU199 cells were routinely maintained in DMEM culture medium supplemented with 10% FBS and 1% Pen-Strep (all from Gibco) and seeded on PLL-coated glass coverslips as needed, allowed to grow overnight and used for experiments thereafter.

### Viruses

We purchased high-titer viruses from BrainVTA (Wuhan) Co. Ltd, viruses were aliquoted and stored at −80°C prior to use. We used an rAAV-9 vector, with a human synapsin (hSyn) promoter, which enabled preferential transduction of neurons. The MscL-G22S sequence was fused with either the fluorescent reporter EYFP or the Ca^2+^ sensor protein GCaMP6S, and a concluding polyA tag. We also used vector controls in addition to the MscL-containing viruses. Viruses used in this study were rAAV/9-hSyn:EYFP-WPRE-pA, rAAV/9-hSyn:MscL-G22S-WPRE-EYFP-pA and rAAV/9- hSyn:MscL-G22S-GCaMP6S-WPRE-pA, rAAV/9-hSyn:GCaMP6S-WPRE-pA.

### Primary cortical neuron culture

Cultured neurons from rat embryos at embryonic day 18 were obtained as previously described^53^. Briefly, cortices were dissected and treated with 0.25% trypsin for 15 min at 37 °C, followed by gentle mixing. The digestion was stopped with Neurobasal medium (Gibco) with 10% fetal bovine serum and 1% penicillin-streptomycin. The cells were resuspended in medium and gently mechanically triturated with a pipette, and then allowed to stand for 15 minutes. The resultant supernatant was discarded, and the cells were resuspended in the abovementioned medium and plated at 1 × 10^5^ cells/cm^2^ in 35 mm dishes with poly-L-lysine-coated (PLL, Gibco) coverslips or PLL-coated glass-bottomed confocal dishes. After 24 hours, the medium was changed to Neurobasal + 2% B27 + 0.25% L-Glutamine + 1% Penicillin-Streptomycin (all from Gibco). Half of the medium was replaced every 2-3 days. Cultured neurons were transducted with AAVs on day 7 and were used in experiments between DIV 10-12 (3-4 days post-infection). All animal studies and experimental procedures were approved by the Animal Subjects Ethics Sub-Committee (ASESC) of the Hong Kong Polytechnic University, and were performed in compliance with the guidelines of the Department of Health - Animals (Control of Experiments) of the Hong Kong S.A.R. government.

### Characterization of ultrasound setup for Ca^2+^ imaging

A flat transducer with center frequency 1.0 MHz (A303S, Olympus) was employed in this study. Ultrasonic pulses were generated using function generator (AFG251, Tektronix) and power amplifier (A075, Electronics & Innovation Ltd.). For ultrasound stimulation, the planar transducer with a diameter of 1.0 cm was fixed perpendicularly downward facing. Cells were grown on glass coverslips, which were held 1.5 cm away from the transducer coupled by plastic wrap encasing degassed deionized water at 25 °C. Acoustic intensity profile was characterized by a hydrophone.

### Cell treatments for calcium imaging

Culture medium was replaced with Fluo-4 AM (5 μM) or X-Rhod-1 AM (10 μM) (both from Invitrogen) working solution in Ca^2+^ solution (pH 7.4), and the cells were incubated at 37 °C in the dark for 30 min. Subsequently, fresh Ca^2+^ solution was used to flush away excess dye before ultrasound stimulation. In mechanistic studies, several different media were used. To remove extracellular Ca^2+^, the coverslip was placed Ca^2+^ free solution with 0.5 mM EGTA to ensure that residual Ca^2+^ was completely chelated. To monitor concurrent cell membrane sonoporation during Ca^2+^ response measurement, the coverslip was perfused with PI solution (100 μg/mL in Ca^2+^ solution, Invitrogen). RR solution (20 μM RR in Ca^2+^ solution, Tocris Bioscience) into the culture medium to evaluate the effect of mechanosensitive ion channels on US+GVs-elicited Ca^2+^ response. 0.20 mM Triton X-100 was added to cells as a positive control of membrane permeability.

### GV-mediated ultrasound stimulation and optical imaging

Briefly, the calcium imaging was done with a modified inverted epifluorescence microscope. The excitation light was generated by a dual-color LED, filtered and delivered to the sample to illuminate the calcium sensor. To minimize phototoxicity, the LEDs were triggered at 1 Hz and synchronized with sCMOS time-lapse imaging. Coverslips with dye-loaded or GCaMP6s-expressing cells were placed above the objective, and GVs were distributed into the media directly before ultrasound stimulation. A camera was used to record the intracellular Fluo-4 AM/X-Rhod-1 AM images with defined time intervals from a function generator at excitation wavelengths of 494 nm for Fluo-4 AM or 580 nm for X-Rhod-1 AM. A bright field image was taken to register the morphology of the cell immediately before and after the GVs mediated ultrasound stimulation. We used software to communicate and coordinate the operation sequence between the microscope and monochromator.

### Evaluation of cytotoxic effects and apoptotic effects

MTT assays was used to evaluate cytotoxicity at different concentration of GVs mediated ultrasound stimulation in the treated CLU199 cells. Cells were treated with GVs alone, or US+GVs in 96-well or 24-well plates. After the indicated treatments and incubations, cells were incubated with 0.5 mg/ml MTT in medium for 3-4 hours at 37°C, solubilized with DMSO and 15 minutes’ shaking, and the solutions’ absorbance at 570 nm was read using an LEDTect 96 microplate reader.

### Western Blot

The treatments’ apoptotic effects were evaluated by a WB of caspase-3. Cells were treated inside an incubator for 15 minutes, allowed to incubate overnight, and protein was collected using RIPA buffer supplemented with 1X Halt Protease and Phosphatase Inhibitor Cocktail (Thermo Scientific). Cells were run on an 4-20% Tris-Glycine SDS-PAGE gel, transferred to activated PVDF membrane (Millipore), and incubated overnight with caspase-3 primary antibody (Cell Signaling #9662) diluted 1:1,000 or α-tubulin primary antibody (Proteintech # 66031-1-Ig) diluted 1:2,500 in 5% milk + TBST. Membranes were washed with TBST, and incubated at room temperature with Goat anti-Rabbit IgG (H+L) superclonal secondary (Invitrogen #A27022) or Rabbit anti-Mouse IgG (H+L) superclonal secondary antibody (Invitrogen #A27033), diluted at 1:10,000 in 5% milk + TBST. Signals were developed using SuperSignal West Pico PLUS Chemiluminescent Substrate and visualized on a ChemiDoc MP imaging system (Bio-Rad). Proteins were quantified using image densitometry and normalized to the α-tubulin expression levels with ImageJ.

### Immunocytochemical fluorescent staining

Cells were treated, allowed to incubate for 90 minutes and fixed using 4% paraformaldehyde + PBS and permeabilized using 0.1% Triton X-100 + PBS, and washes were done with 1X PBS or 1X PBS+Tween-20 (PBST) (after permeabilization). Cells were blocked with 2% BSA + 0.3M Glycine + PBST, and incubated overnight with primary antibodies in 2% BSA + PBST. The next day, cells were washed and incubated with secondary antibodies in 2% BSA + PBST, then washed and mounted with Fluoroshield Mounting Medium with DAPI (Abcam). Stained cells were imaged on a Leica TCS SP8 confocal microscope. All steps from secondary antibody incubation onwards were performed in the dark.

Primary antibodies used were c-Fos (Cell Signaling #2250) at a dilution of 1: 3,000, and MAP2 (PA1-10005, Invitrogen) at a dilution of 1:2,500. Secondary antibodies, used at a dilution of 1:1,000, were Goat anti-Rabbit IgG Alexa Fluor Plus 488 (#A32731), Goat anti-Chicken IgY Alexa Fluor Plus 555(#A32932) or Goat anti-Rabbit IgG Alexa Fluor Plus 555 (#A32732), all from Invitrogen.

### c-Fos counting

The number of c-Fos^+^ cells in primary neurons was determined by counting the number of neuronal nuclei showing c-Fos expression 90 minutes after stimulation. For non-transducted cells, nuclear c-Fos was counted in cells staining positive for MAP2. For transducted cells, nuclear c-Fos was counted in cells showing EYFP expression. The percentage of cells showing c-Fos among the cells identified was then calculated per experiment. We also calculated the number of EYFP^+^ and EYFP^-^ cells with nuclear c-Fos expression in MscL-transducted dishes. Each experiment had a minimum of 10 photographed FOVs and minimum of 50 total cells counted per condition.

### Statistical analyses

A minimum of 3 independent experiments were performed for all experiments shown, meaning at least 3 separate ‘rounds’ of cell preparations, transfections or primary neuron harvests that were used for various experiments. Wherever possible, multiple plates from each round were evaluated. The data were collected into GraphPad Prism sheets for statistical analysis and graph preparation. Two-tailed unpaired *t*-tests or one-way or two-way ANOVA were performed to determine statistical significance, with post-hoc tests or corrections applied where appropriate. P values below 0.05 were considered significant.

## Supporting information

**Supplementary Fig. 1.**
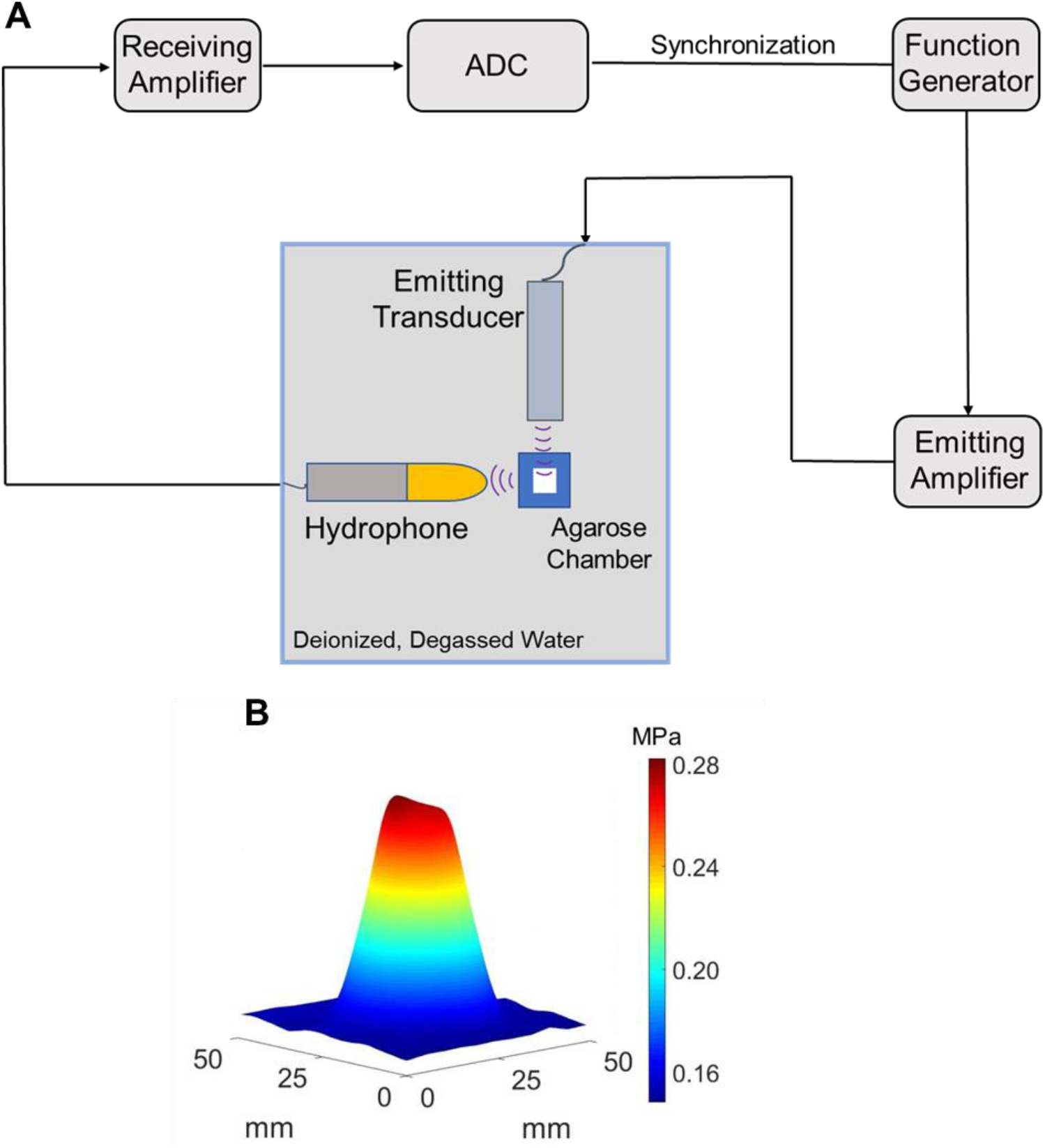
Cavitation detection system and acoustic field characterization. **a,** The *in vitro* passive cavitation detection system used to measure the backscattered acoustic signals of cavitation in response to GVs + ultrasound stimulation. **b,** Acoustic field characterization of our stimulation setup, with a spatial PNP of 0.28 MPa.

**Supplementary Fig. 2.**
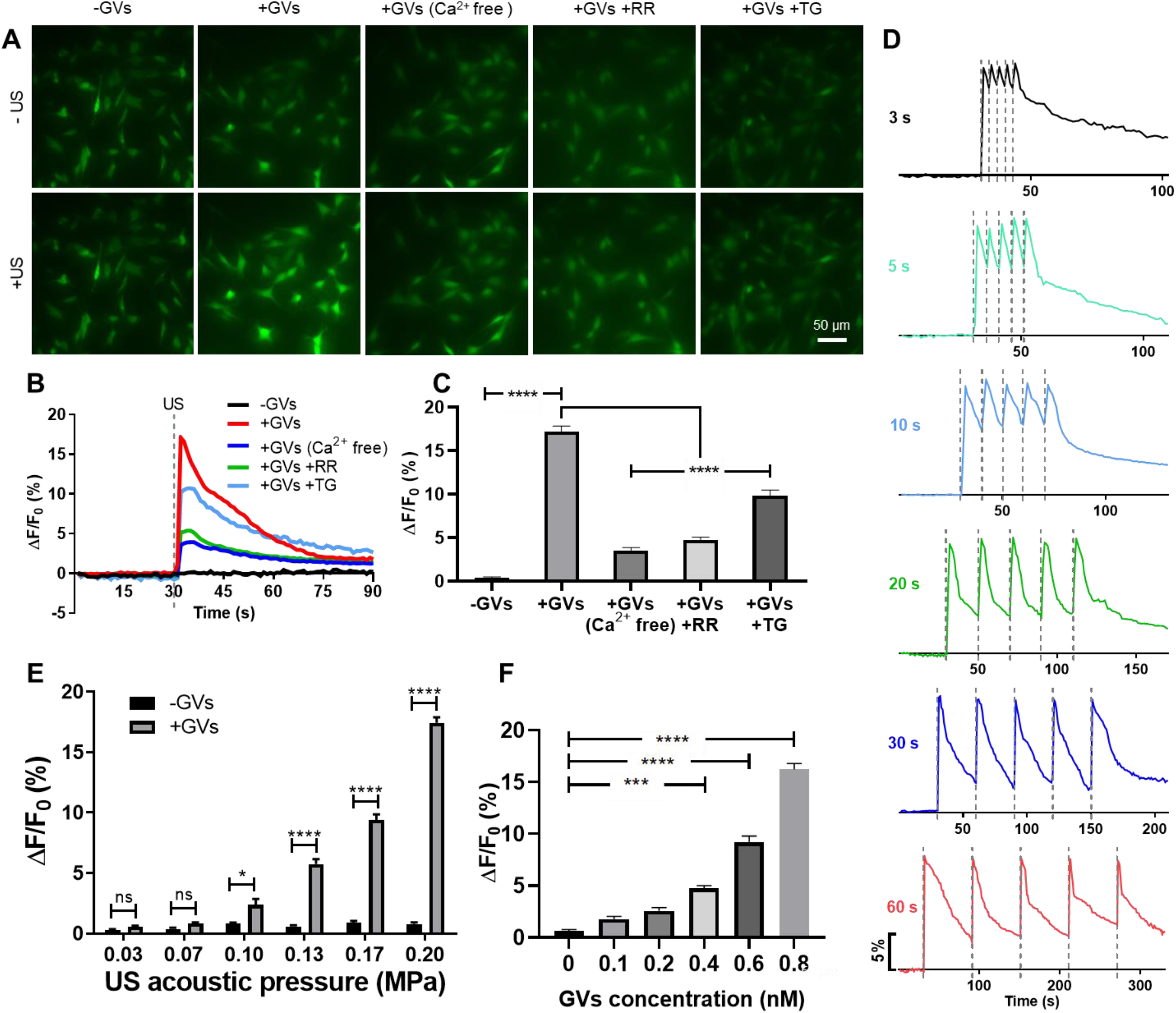
Calcium imaging of ultrasound + GVs stimulation performed in the neuronal cell line, CLU199. **a,** Representative images of the typical Ca^2+^ response seen in CLU199 cells before and after 0.20 MPa ultrasound stimulation with or without GVs, in calcium-free medium, with the broad-spectrum mechanosensitive ion channel blocker ruthenium red (RR) or with the internal calcium chelator Thapsigargin (TG). **b,** Time-course of the imaging results depicted in **(a)**. **c,** Quantification of the fluorescence intensity changes shown in **(a)** and **(b)**. Bars represent mean ± SEM of 3 independent experiments. ****, p < 0.0001 compared only to the +GVs condition, unpaired one-way ANOVA with post-hoc Dunnett test. **d,** Time-resolved Ca^2+^ responses of CLU199 cells stimulated by 5 ultrasound pulses at varying intervals. **e,** Ca^2+^ response of cells to varying ultrasound intensities, 0.8 nM GV. Bars represent the mean ± SEM of 3 independent experiments. *, p < 0.05; ****, p < 0.0001; two-way ANOVA with Sidak correction. **f,** Ca^2+^ response of cells to varying GVs concentrations, 0.20 MPa ultrasound. Bars represent the mean ± SEM of 3 independent experiments. *, p < 0.05; p < 0.0001 compared only to the 0 nM GVs condition, unpaired one-way ANOVA with Dunnett correction.

**Supplementary Fig. 3.**
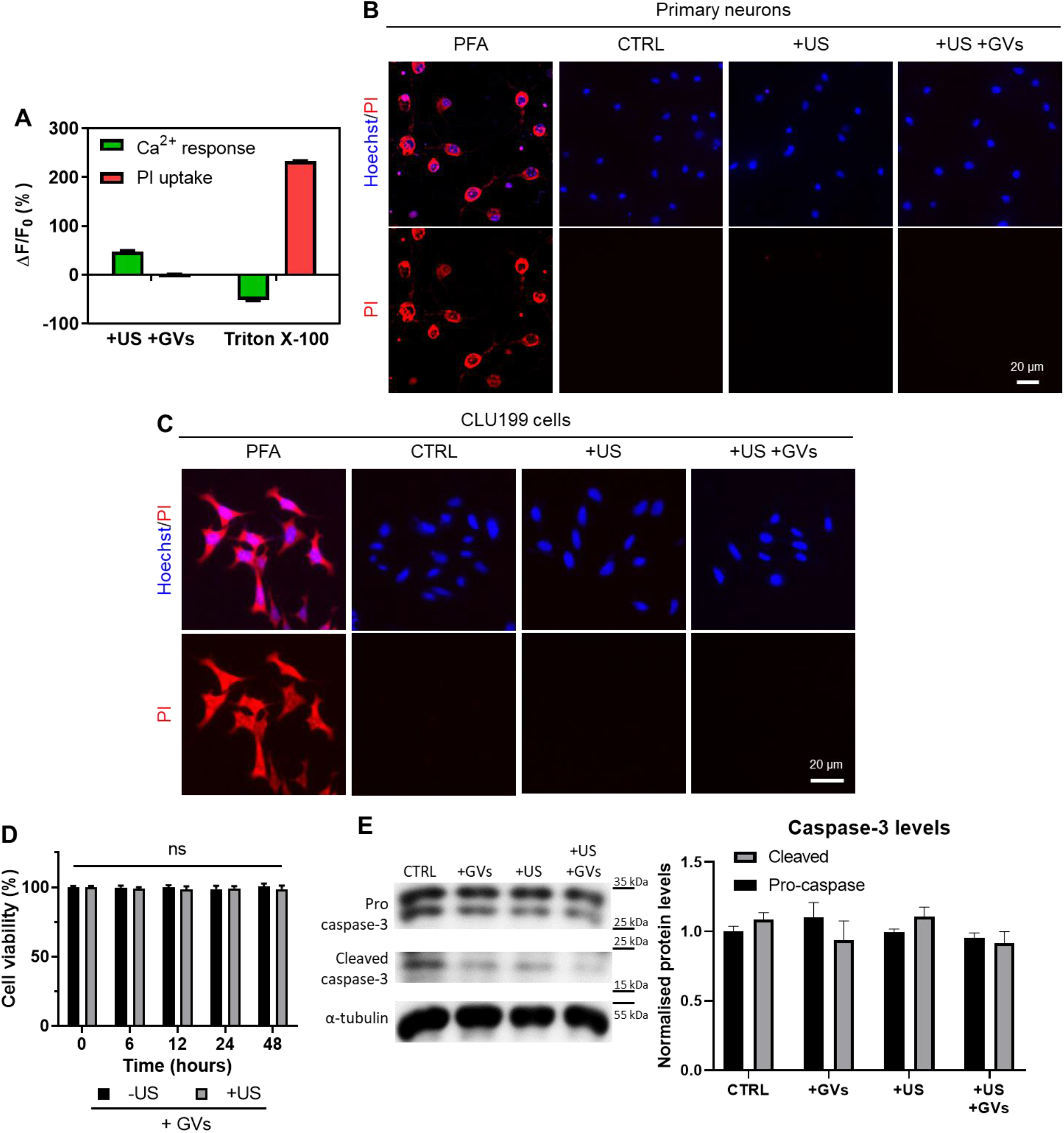
Evidence for the non-cytotoxicity of our GV-mediated ultrasound treatment. **a,** Quantified fluorescence changes of neuronal Ca^2+^ response and propidium iodide (PI) uptake of the representative images shown in Fig. 3A. Bars represent mean ± SD from 3 independent experiments. **b,** Intracellular uptake of PI by primary neurons when untreated, treated with ultrasound-alone or with ultrasound + GVs (0.20 MPa, 10 second interval, 10% duty cycle, 0.8 nM GVs). Primary neurons treated with 4% paraformaldehyde (PFA) are shown here as a positive control for membrane permeation and PI staining. **c,** Intracellular uptake of PI by CLU199 cells. All treatment conditions were the same as in **(b). d,** Cell viability following US+GVs treatments. CLU199 cells were treated with either GVs alone or US+GVs for 15 minutes, and their cell viability at various times post-treatment was determined using an MTT assay. Bars represent the mean ± SEM of 3 independent experiments. No significant differences found, multiple two-tailed *t*-tests with Holm-Sidak correction. **e,** Caspase-3 levels in primary neurons following various treatments. Primary neurons were exposed to either GVs, ultrasound or ultrasound + GVs for 15 minutes, proteins were collected after overnight incubation and a WB was performed to observe levels of pro-and cleaved caspase-3. Only upper bands were quantified for pro caspase-3. Bars represent the mean ± SEM of 3 independent experiments. No significant differences found, two-way ANOVA with post-hoc Tukey test.

**Supplementary Fig. 4.**
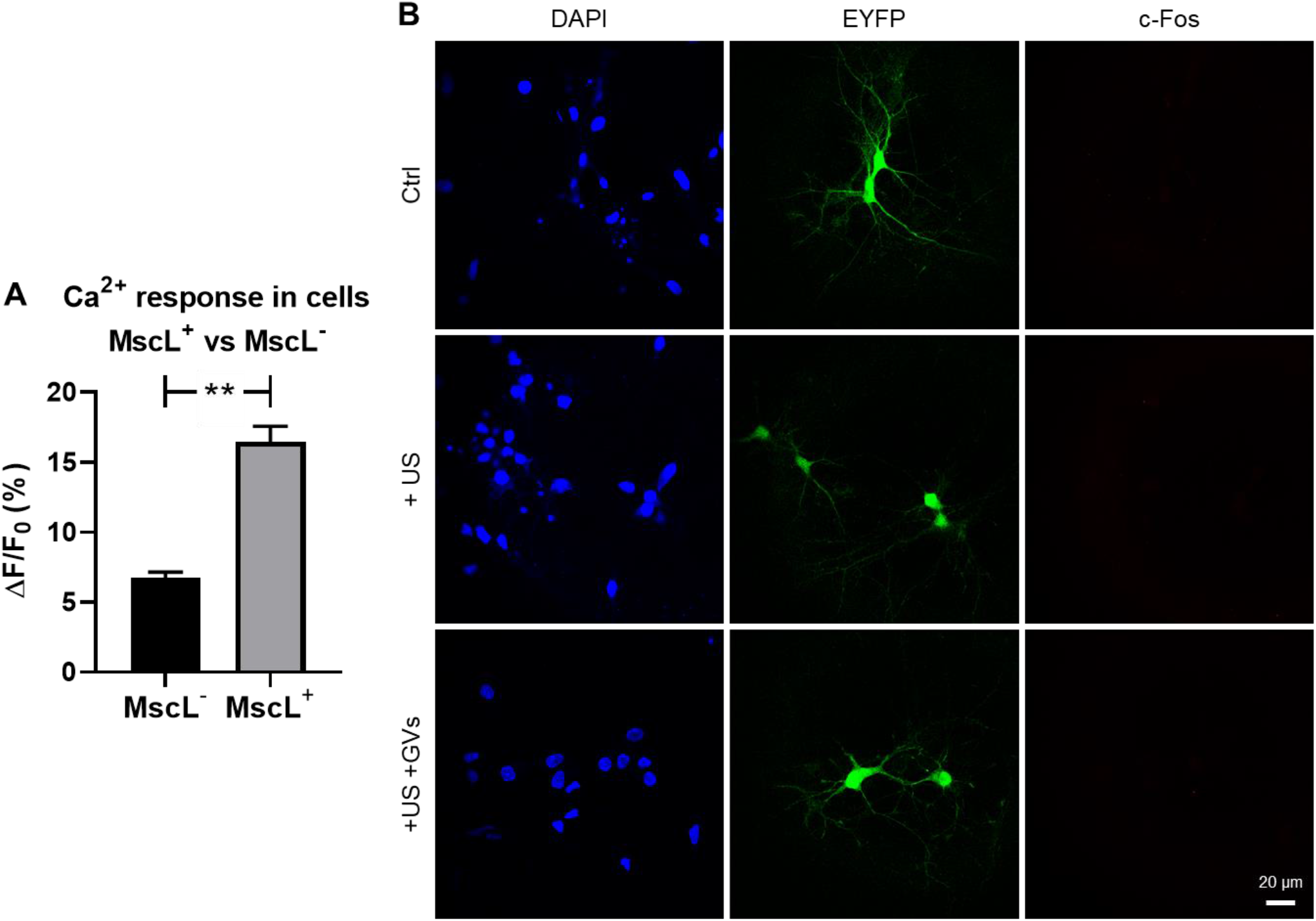
Further Ca^2+^ imaging and c-Fos staining data from transducted neurons. **a**, Quantified fluorescence intensity changes in primary neurons in a dish transducted with MscL-EYFP virus, as shown in Supplementary Video 3. Ca^2+^ intensities of cells in the same dish were quantified and grouped according to whether the cells expressed EYFP (‘MscL^+^’) or did not express EYFP (‘MscL^-^’). **, p < 0.01, two-tailed unpaired *t*-test. **b**, Representative images of neuronal c-Fos expression in cells transducted with EYFP AAVs in untreated cells, or cells treated with ultrasound alone or US+GVs.

### Figure Legends for Supplementary Videos 1 – 3

**Video 1.** Primary neurons show rapid and reversible calcium influx in response to each ultrasound pulse (0.20 MPa) when GVs (0.8 nM) are present (8 pulses delivered).

**Video 2.** GV-mediated ultrasound stimulation (0.20 MPa) triggers calcium influx into primary neurons, without allowing PI to enter cells (showing harmful membrane permeation).

**Video 3.** Primary neurons expressing the mechanosensitive ion channel MscL-G22S-EYFP display stronger responses to US+GVs (0.13 MPa, 0.4 nM) than EYFP^-^ cells.

## Notes

### Competing Interest Statement

The authors have declared no competing interest.

### Summary of Updates

The tittle has been updated.

## References

1. Chen S, Weitemier AZ, Zeng X, He L, Wang X, Tao Y, et al. Near-infrared deep brain stimulation via upconversion nanoparticle-mediated optogenetics. Science 2018, 359(6376): 679–684.

2. Carvalho-de-Souza JL, Treger JS, Dang B, Kent SB, Pepperberg DR, Bezanilla F. Photosensitivity of neurons enabled by cell-targeted gold nanoparticles. Neuron 2015, 86(1): 207–217.

3. Chen R, Romero G, Christiansen MG, Mohr A, Anikeeva P. Wireless magnetothermal deep brain stimulation. Science 2015, 347(6229): 1477–1480.

4. Tay A, Di Carlo D. Magnetic Nanoparticle-Based Mechanical Stimulation for Restoration of Mechano-Sensitive Ion Channel Equilibrium in Neural Networks. Nano Lett 2017, 17(2): 886–892.

5. Gregurec D, Senko AW, Chuvilin A, Reddy PD, Sankararaman A, Rosenfeld D, et al. Magnetic Vortex Nanodiscs Enable Remote Magnetomechanical Neural Stimulation. ACS Nano 2020, 14(7): 8036–8045.

6. Tyler WJ, Lani SW, Hwang GM. Ultrasonic modulation of neural circuit activity. Curr Opin Neurobiol 2018, 50: 222–231.

7. Tufail Y, Yoshihiro A, Pati S, Li MM, Tyler WJ. Ultrasonic neuromodulation by brain stimulation with transcranial ultrasound. Nat Protoc 2011, 6(9): 1453–1470.

8. Li G, Qiu W, Zhang Z, Jiang Q, Su M, Cai R, et al. Noninvasive ultrasonic neuromodulation in freely moving mice. 2018, 66(1): 217–224.

9. Yoo S-S, Bystritsky A, Lee J-H, Zhang Y, Fischer K, Min B-K, et al. Focused ultrasound modulates region-specific brain activity. 2011, 56(3): 1267–1275.

10. Dallapiazza RF, Timbie KF, Holmberg S, Gatesman J, Lopes MB, Price RJ, et al. Noninvasive neuromodulation and thalamic mapping with low-intensity focused ultrasound. J Neurosurg 2018, 128(3): 875–884.

11. Lee W, Lee SD, Park MY, Foley L, Purcell-Estabrook E, Kim H, et al. Image-guided focused ultrasound-mediated regional brain stimulation in sheep. 2016, 42(2): 459–470.

12. Verhagen L, Gallea C, Folloni D, Constans C, Jensen DE, Ahnine H, et al. Offline impact of transcranial focused ultrasound on cortical activation in primates. 2019, 8: e40541.

13. Legon W, Ai L, Bansal P, Mueller JK. Neuromodulation with single-element transcranial focused ultrasound in human thalamus. Hum Brain Mapp 2018, 39(5): 1995–2006.

14. Lee W, Kim HC, Jung Y, Chung YA, Song IU, Lee JH, et al. Transcranial focused ultrasound stimulation of human primary visual cortex. Sci Rep 2016, 6: 34026.

15. Legon W, Bansal P, Tyshynsky R, Ai L, Mueller JK. Transcranial focused ultrasound neuromodulation of the human primary motor cortex. Sci Rep 2018, 8(1): 10007.

16. Legon W, Sato TF, Opitz A, Mueller J, Barbour A, Williams A, et al. Transcranial focused ultrasound modulates the activity of primary somatosensory cortex in humans. Nature Neuroscience 2014, 17(2): 322–329.

17. Lee W, Kim H, Jung Y, Song IU, Chung YA, Yoo SS. Image-guided transcranial focused ultrasound stimulates human primary somatosensory cortex. Sci Rep 2015, 5: 8743.

18. Leo A, Mueller JK, Grant A, Eryaman Y, Wynn L. Transcranial focused ultrasound for BOLD fMRI signal modulation in humans. Conf Proc IEEE Eng Med Biol Soc 2016, 2016: 1758–1761.

19. Mueller J, Legon W, Opitz A, Sato TF, Tyler WJ. Transcranial focused ultrasound modulates intrinsic and evoked EEG dynamics. Brain Stimul 2014, 7(6): 900–908.

20. Beisteiner R, Matt E, Fan C, Baldysiak H, Schonfeld M, Philippi Novak T, et al. Transcranial Pulse Stimulation with Ultrasound in Alzheimer’s Disease-A New Navigated Focal Brain Therapy. Adv Sci (Weinh) 2020, 7(3): 1902583.

21. Meng Y, Volpini M, Black S, Lozano AM, Hynynen K, Lipsman N. Focused Ultrasound as a Novel Strategy for Alzheimer Disease Therapeutics. Ann Neurol 2017, 81(5): 611–617.

22. Lin Z, Meng L, Zou J, Zhou W, Huang X, Xue S, et al. Non-invasive ultrasonic neuromodulation of neuronal excitability for treatment of epilepsy. Theranostics 2020, 10(12): 5514–5526.

23. Tsai SJ. Transcranial focused ultrasound as a possible treatment for major depression. Med Hypotheses 2015, 84(4): 381–383.

24. Abrahao A, Meng Y, Llinas M, Huang Y, Hamani C, Mainprize T, et al. First-in-human trial of blood–brain barrier opening in amyotrophic lateral sclerosis using MR-guided focused ultrasound. 2019, 10(1): 1–9.

25. Kubanek J, Shi J, Marsh J, Chen D, Deng C, Cui J. Ultrasound modulates ion channel currents. Sci Rep 2016, 6: 24170.

26. Kubanek J, Shukla P, Das A, Baccus SA, Goodman MB. Ultrasound Elicits Behavioral Responses through Mechanical Effects on Neurons and Ion Channels in a Simple Nervous System. Journal of Neuroscience 2018, 38(12): 3081–3091.

27. Tyler WJ. The mechanobiology of brain function. Nat Rev Neurosci 2012, 13(12): 867–878.

28. Bourdeau RW, Lee-Gosselin A, Lakshmanan A, Farhadi A, Kumar SR, Nety SP, et al. Acoustic reporter genes for noninvasive imaging of microorganisms in mammalian hosts. 2018, 553(7686): 86–90.

29. Yang Y, Qiu Z, Hou X, Sun LJUim, biology. Ultrasonic Characteristics and Cellular Properties of Anabaena Gas Vesicles. 2017, 43(12): 2862–2870.

30. Buckland B, Walsby AJAfM. A study of the strength and stability of gas vesicles isolated from a blue-green alga. 1971, 79(4): 327–337.

31. Honary S, Zahir F. Effect of Zeta Potential on the Properties of Nano-Drug Delivery Systems - A Review (Part 2). Trop J Pharm Res 2013, 12(2): 265–273.

32. Coussios CC, Roy RA. Applications of acoustics and cavitation to noninvasive therapy and drug delivery. Annu Rev Fluid Mech 2008, 40: 395–420.

33. Qiu Z, Guo J, Kala S, Zhu J, Xian Q, Qiu W, et al. The Mechanosensitive Ion Channel Piezo1 Significantly Mediates In Vitro Ultrasonic Stimulation of Neurons. iScience 2019, 21: 448–457.

34. Chaudhuri A, Zangenehpour S, Rahbar-Dehgan F, Ye FC. Molecular maps of neural activity and quiescence. Acta Neurobiol Exp 2000, 60(3): 403–410.

35. Koley D, Bard AJ. Triton X-100 concentration effects on membrane permeability of a single HeLa cell by scanning electrochemical microscopy (SECM). Proc Natl Acad Sci U S A 2010, 107(39): 16783–16787.

36. Li F, Yang C, Yuan F, Liao D, Li T, Guilak F, et al. Dynamics and mechanisms of intracellular calcium waves elicited by tandem bubble-induced jetting flow. Proc Natl Acad Sci U S A 2018, 115(3): E353–E362.

37. !!! INVALID CITATION !!!.

38. Qiu Z, Kala S, Guo J, Xian Q, Zhu J, Zhu T, et al. Targeted Neurostimulation in Mouse Brains with Non-invasive Ultrasound. Cell Rep 2020, 32(7): 108033.

39. Bunney PE, Zink AN, Holm AA, Billington CJ, Kotz CM. Orexin activation counteracts decreases in nonexercise activity thermogenesis (NEAT) caused by high-fat diet. Physiol Behav 2017, 176: 139–148.

40. Tyler WJ, Tufail Y, Finsterwald M, Tauchmann ML, Olson EJ, Majestic C. Remote Excitation of Neuronal Circuits Using Low-Intensity, Low-Frequency Ultrasound. Plos One 2008, 3(10).

41. Segel M, Neumann B, Hill MFE, Weber IP, Viscomi C, Zhao C, et al. Niche stiffness underlies the ageing of central nervous system progenitor cells. Nature 2019, 573(7772): 130–134.

42. Zhang J, Zhou Y, Huang T, Wu F, Liu L, Kwan JSH, et al. PIEZO1 functions as a potential oncogene by promoting cell proliferation and migration in gastric carcinogenesis. Molecular Carcinogenesis 2018, 57(9): 1144–1155.

43. Aykut B, Chen R, Kim JI, Wu D, Shadaloey SAA, Abengozar R, et al. Targeting Piezo1 unleashes innate immunity against cancer and infectious disease. Sci Immunol 2020, 5(50).

44. Wang GH, Song L, Hou XD, Kala S, Wong KF, Tang LY, et al. Surface-modified GVs as nanosized contrast agents for molecular ultrasound imaging of tumor. Biomaterials 2020, 236.

45. Farhadi A, Ho GH, Sawyer DP, Bourdeau RW, Shapiro MG. Ultrasound imaging of gene expression in mammalian cells. Science 2019, 365(6460): 1469–+.

46. Alcaino C, Farrugia G, Beyder A. Mechanosensitive Piezo Channels in the Gastrointestinal Tract. Curr Top Membr 2017, 79: 219–244.

47. Dong Q, He L, Chen LB, Deng QZ. Opening the Blood-Brain Barrier and Improving the Efficacy of Temozolomide Treatments of Glioblastoma Using Pulsed, Focused Ultrasound with a Microbubble Contrast Agent. Biomed Res Int 2018, 2018.

48. Song KH, Fan AC, Hinkle JJ, Newman J, Borden MA, Harvey BK. Microbubble gas volume: A unifying dose parameter in blood-brain barrier opening by focused ultrasound. Theranostics 2017, 7(1): 144–152.

49. Song KH, Harvey BK, Borden MA. State-of-the-art of microbubble-assisted blood-brain barrier disruption. Theranostics 2018, 8(16): 4393–4408.

50. Tsai HC, Tsai CH, Chen WS, Inserra C, Wei KC, Liu HL. Safety evaluation of frequent application of microbubble-enhanced focused ultrasound blood-brain-barrier opening. Sci Rep-Uk 2018, 8.

51. Quaia E. Microbubble ultrasound contrast agents: an update. Eur Radiol 2007, 17(8): 1995–2008.

52. Yang HL, Cai WB, Xu L, Lv XH, Qiao YB, Li P, et al. Nanobubble-Affibody: Novel ultrasound contrast agents for targeted molecular ultrasound imaging of tumor. Biomaterials 2015, 37: 279–288.

53. Pi RB, Li WM, Lee NTK, Chan HHN, Pu YM, Chan LN, et al. Minocycline prevents glutamate-induced apoptosis of cerebellar granule neurons by differential regulation of p38 and Akt pathways. J Neurochem 2004, 91(5): 1219–1230.

